# RemoteFoldSet: Benchmarking Structural Awareness of Protein Language Models

**DOI:** 10.1101/2025.09.23.678152

**Authors:** Zinnia Ma, Neville P. Bethel

## Abstract

Protein language models (pLMs) have the capacity to infer structural information from amino acid sequences. Evaluating the extent to which structural information they truly encode is crucial for assessing their generalizability and the interpretability of their latent representations, yet current approaches lack a model-free, quantitative framework to evaluate these encodings. We introduce RemoteFoldSet, a curated collection of protein sequence sets stratified by high structural similarity but minimal sequence identity. We also define the Structural Awareness (SA) score, a novel metric that enables model-agnostic, unsupervised, and training-free quantification of structure-related patterns in pLM embeddings. Using RemoteFoldSet together with the SA score, we benchmark a range of existing pLMs, elucidating how models with different training objectives, architectures, and sizes discriminate and distribute proteins within their embedding spaces, both quantitatively and qualitatively. We expect that this methodology will serve as a reliable benchmark for evaluating the performance of pLMs for structural and functional applications.

## 1 Introduction

As a paradigm of applying natural language processing (NLP) techniques to protein sequences, pLMs have become indispensable in drug design and computational biology, facilitating both the prediction of protein structure and protein complexes as well as the generative design of *de novo* proteins. The goal of a pLM is to extend sequence-level patterns to a protein’s structure and, ultimately, its function. How a pLM organizes its embedding space is therefore critical to making these connections. Assessing the extent to which pLMs capture structural information is essential not only for ensuring robust generalization, but also for interpreting their internal representations in biologically meaningful ways. For example, in protein–protein interaction prediction, a model with structural awareness may generalize well by learning the underlying physicochemical principles that confer specificity and affinity. By contrast, a model lacking structural awareness might still achieve high accuracy for benchmark datasets by memorizing superficial patterns at the sequence level, but such reliance can undermine robustness and lead to failures in novel or distribution-shifted scenarios[1]. Thus, we must understand the structure of the pLM embedding space to analyze whether the pLM explicitly encodes true physicochemical features or merely captures spurious patterns.

Structure prediction models like ESMfold utilize pLM embeddings to predict protein structure in a supervised manner [2], but pLMs have been shown to be capable of inferring structural information from sequence alone. In many transformer-based pLMs, researchers have observed that attention patterns between residues often correlate with their spatial interactions[3][4]. Other studies applying linear probes to assess pLM-derived embeddings have demonstrated that these embeddings can directly predict secondary structures and contact maps[5]. These results suggest that these representations indeed capture meaningful structural information without explicitly being trained on structural data. The pLM research space has been expanding rapidly with a wide range in model sizes, layers, architectures and training sets. The extent to which these models capture structural features is unclear, and to date, no model-free approach currently exists to quantitatively evaluate the degree to which structural information is captured in these representations.

To address this evaluation gap, we propose a benchmark dataset specifically designed to decouple structural homology from sequence identity. Each set in the dataset consists of protein sequences with extremely low sequence identity but highly consistent 3D structures. We introduce a metric that probes the organization of the pLM embedding space, investigating if protein sequences of similar structure, but low identity, cluster in the embedding space. By computing pairwise similarities between the sequence embeddings produced by different pLMs within each set, we quantify this clustering and compare these pairwise similarities across pLM models. Because sequence identity is minimized, any similarity in the embeddings can be attributed to shared structural features rather than sequence-level patterns.

In this study, we investigate the structural awareness encoded in pLMs by proposing a novel model-agnostic and unsupervised evaluation framework. Our contributions are summarized as follows:

- **Construction of RemoteFoldSet**: a new dataset comprising 90 protein sets of diverse folds with alternative, low-identity sequences generated via ProteinMPNN[6] and filtered for foldability using AlphaFold3[7]. Each set includes sequence-diverse proteins with consistent folds, enabling evaluation decoupled from sequence similarity.
- **Structure-aware evaluation method**: a new metric to assess pLM embeddings that is model-agnostic, unsupervised, and training-free, and supports layer-wise analysis of structural information encoded across model depths.
- **Extensive benchmarking of pLMs**: applying our framework to widely-used pLMs reveals differences in their capacity to capture structural similarity, offering insight into their representational strengths and limitations.

## 2 Related Work

Traditionally, pLMs have been evaluated through supervised downstream tasks. The TAPE benchmark [5] introduced a suite of biologically relevant, semi-supervised tasks to assess protein embeddings, including secondary structure prediction and contact map prediction, two structurally informative tasks. However, because these tasks rely on labeled data and fine-tuning, they may not directly reflect the intrinsic structural awareness encoded in pLM representations.

More recent benchmarks, such as FLIP[8] and ProteinGym[9], focus on mutation effect prediction and fitness estimation. These benchmarks typically compare models on their ability to predict outcomes of experimental assays that map protein sequences to phenotypic measurements. While such tasks may correlate with structural properties, they are primarily designed to evaluate functional fitness rather than the quality of structural representations.

In contrast, our proposed RemoteFoldSet provides a model-agnostic and unsupervised framework for evaluating structural awareness. It operates purely on sequence embeddings, leveraging structure-consistent but sequence-divergent protein sets to assess whether pLMs capture fold-level similarity without relying on downstream tasks or structural supervision.

## 3 Methods

### 3.1 Dataset generation

To ensure that the selected starting structures are distributed across the known structural space, we sample proteins from clusters of the CATH dataset[10][11], a widely used resource that offers a hierarchical classification of protein domain structures that sorts proteins by class, architecture, fold, superfamily, and then domain. We selected domains that cover all 40 architectures within the three primary CATH classes: Mainly Alpha, Mainly Beta, and Alpha Beta. To minimize redundancy and ensure structural diversity, no two domains were selected from the same superfamily. These 168 starting structures were used as the input backbones for alternative sequence generation.

Constructing such a dataset while minimizing sequence similarity and preserving structural consistency requires a mechanism to generate sequence-diverse proteins with shared structures. To this end, we leverage inverse folding models, which predict amino acid sequences that likely fold into a given protein’s three-dimensional structure. We employ ProteinMPNN, which is a message passing neural network capable of stochastically decoding protein sequences using a Monte Carlo, temperature dependent, protocol. Starting from a fixed structure, we generate diverse candidate sequences by raising the temperature of the sampling module, which encourages exploration of a broader sequence space. We set the temperature to T=1.0 and generate 160 sequences per structure.

Mutations in a protein sequence will inevitably alter a protein’s ground state structure, but some mutations or alternative sequences will fold into a very similar structure while others may either not fold or adopt a different conformation. To ensure that these sampled sequences indeed correspond to a similar structure, we validate each one using Alphafold3. Since our primary interest was in the overall protein fold, we used the pTM score to assess the reliability of the structures predicted by AlphaFold3, and the TM-score to evaluate their agreement with the given structures. From the 160 generated sequences of each domain, we retained only those satisfying both pTM > 0.8 and TM-score > 0.8, resulting in 9,893 out of 26,880 sequences across 100 domains (Table 3). Then a greedy search was applied to select the 16 most diverse sequences. Domains with fewer than 16 sequences that met the criteria were discarded. This process yielded a dataset of 90 protein sets, each containing 16 sequences (1,440 sequences in total), with a mean sequence diversity of 0.74.

Although our dataset does not yet reach the ideal scenario of near-zero sequence identity, the mean sequence identity is approximately 26%, placing it squarely in the so-called ‘twilight zone’ of protein sequence similarity, where homology detection becomes unreliable without structural information[12]. By having sequences with similar fold but almost no identity, we can evaluate the embedding space of these proteins and determine if the embedding space will distribute the proteins according to their similar structure rather than sequence. This will give us insight into whether these pLMs have ‘structural awareness’ from sequence alone.

### 3.2 Detect structural awareness

With the validated dataset, we evaluated the extent to which the embeddings derived from pLMs contain structural information. All 1,440 sequences in the dataset were fed into the pLMs (Table 4) to obtain their corresponding embeddings. To remove global bias, we applied mean centering to all embeddings from each model, i.e., subtracting the mean embedding vector from each individual embedding.

For each structural set 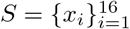, the model *f* generates sequence embeddings *z*_*i*_ = *f* (*x*_*i*_). After mean-centering all *{z*_*i*_*}*, we compute the pairwise cosine similarities and take their average as the **Structural Awareness (SA)** score for that set:

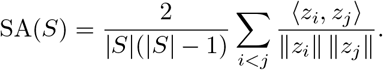

This metric quantifies the extent to which embeddings capture fold-level similarity beyond sequence identity. Unless otherwise stated, we report the mean and standard deviation (mean ± std) of SA scores between different sets. Since the 90 protein sets correspond to domains distributed across the structural space, we further group them by their CATH class-level annotations to enable a more comprehensive evaluation. Proteins from different structural classes often exhibit distinct folding patterns, so analyzing model performance within each group provides additional insight into the structural sensitivity of pLMs. To validate the robustness of our metric, we also compute SA scores on randomly shuffled embeddings as a baseline control (Figure 2).

**Figure 1:**
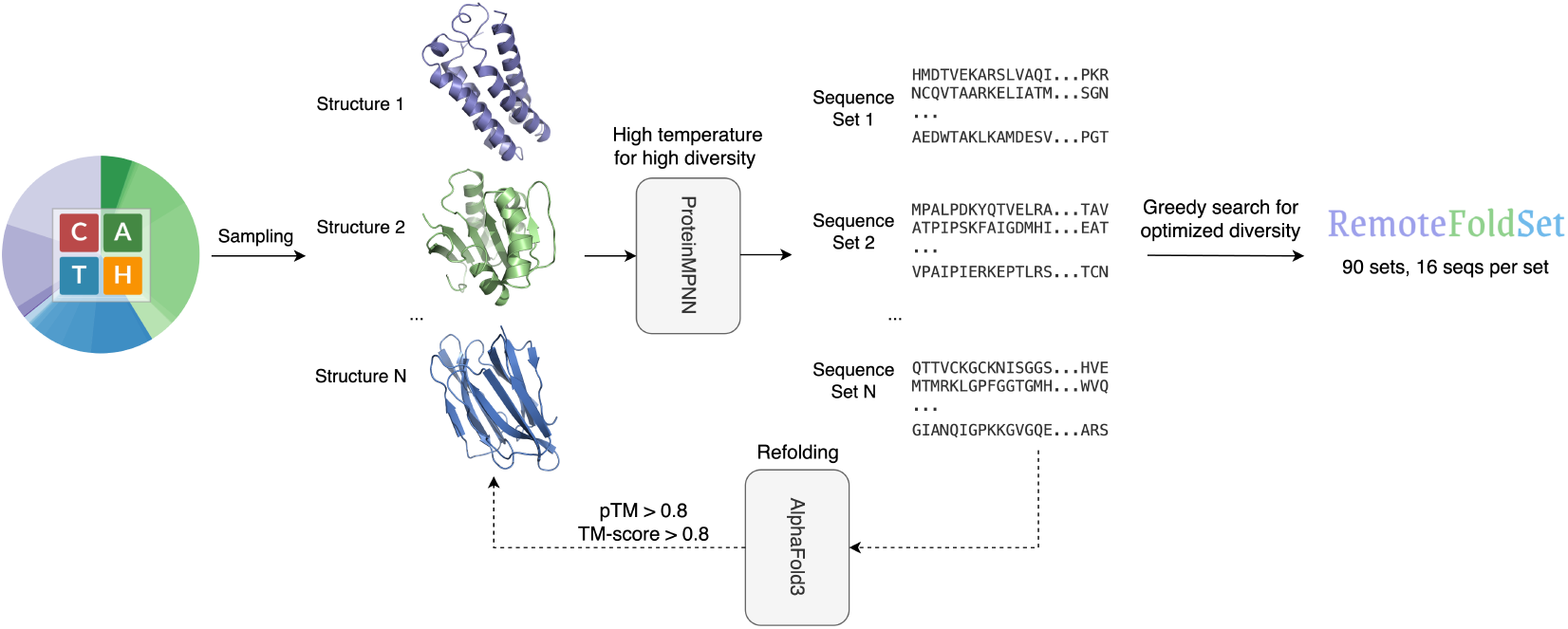
RemoteFoldSet workflow. Protein domains are selected from CATH, sequence variants are generated via ProteinMPNN, filtered using AlphaFold3, and grouped into structure-consistent, sequence-diverse sets.

**Figure 2:**
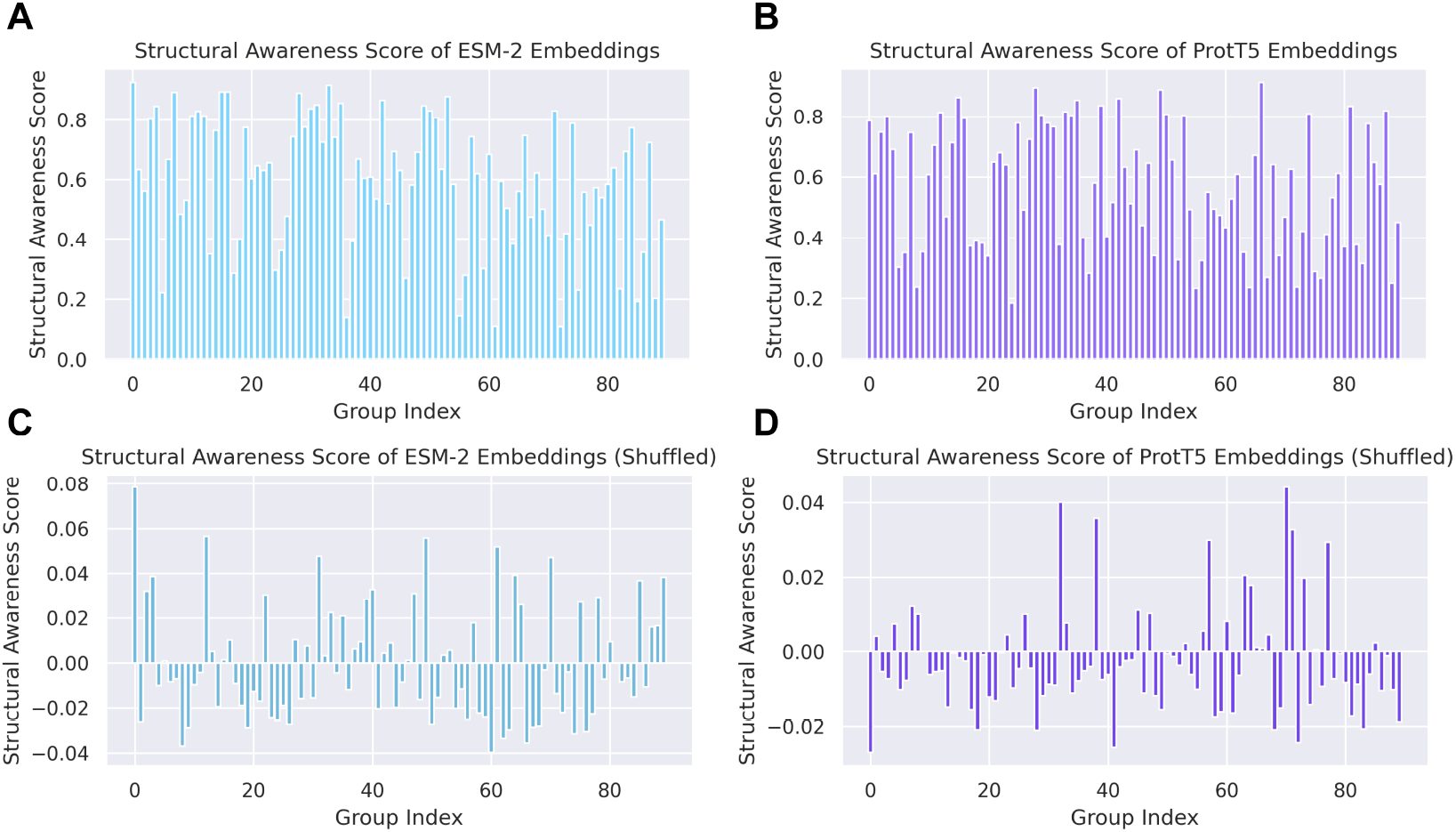
Structural awareness score of pLM embeddings and shuffled embeddings. **(A-B)** SA score of the 90 embedding groups according to the 90 protein sets. **(C-D)** SA score of the 90 shuffled embedding groups.

While the SA score quantifies the overall amount of structural information by measuring fold-level similarity, the quantity of information is not always equivalent to its quality. For example, certain models may achieve high SA scores by capturing shared local structural patterns influenced by architectural inductive biases, which may not support reliable distinctions between different folds. To mitigate this, we additionally define the SA distance ratio, which compares intra-group and inter-group distances as a simple auxiliary metric for assessing the discriminability of encoded structural information. A lower ratio suggests better structural discrimination beyond global similarity (Appendix A.3).

## 4 Experiments

### 4.1 Benchmarking protein language models

We first benchmark multiple pLMs on the RemoteFoldSet dataset using the proposed SA metric. This benchmark quantifies how well each model captures fold-level structure beyond sequence similarity. For the benchmark, we include a representative selection of pLMs, comprising UniRep[13], multiple variants of ProGen2[14], CARP[15], several models from the ESM family[2], as well as the ProtT5 and ProtBert models from the ProtTrans suite[4]. The detailed results are presented in Table 1.

**Table 1:** Comparison of the structural awareness scores of different pLMs (higher is better).

| Models | All domains | Mainly Alpha | Mainly Beta | Alpha Beta |
| --- | --- | --- | --- | --- |
| UniRep | 0.52 ± 0.19 | **0.64 ± 0.10** | 0.54 ± 0.21 | 0.45 ± 0.17 |
| ProGen2 (base) | 0.45 ± 0.21 | 0.46 ± 0.17 | 0.51 ± 0.23 | 0.36 ± 0.17 |
| ProGen2 (large) | 0.45 ± 0.18 | 0.51 ± 0.11 | 0.49 ± 0.21 | 0.38 ± 0.11 |
| ProGen2 (xlarge) | 0.33 ± 0.25 | 0.43 ± 0.28 | 0.37 ± 0.25 | 0.25 ± 0.19 |
| CARP (640M) | 0.61 ± 0.28 | 0.59 ± 0.12 | **0.70 ± 0.27** | 0.50 ± 0.30 |
| ESM-1b | 0.59 ± 0.21 | 0.54 ± 0.17 | 0.65 ± 0.21 | 0.54 ± 0.21 |
| ESM-2 (650M) | 0.59 ± 0.22 | 0.51 ± 0.17 | 0.65 ± 0.21 | 0.55 ± 0.22 |
| ESM-2 (3B) | **0.63 ± 0.20** | 0.57 ± 0.16 | 0.67 ± 0.20 | **0.60 ± 0.20** |
| ESM-2 (15B) | 0.60 ± 0.19 | 0.54 ± 0.17 | 0.65 ± 0.19 | 0.57 ± 0.18 |
| ProtBert | 0.54 ± 0.20 | 0.59 ± 0.09 | 0.57 ± 0.22 | 0.49 ± 0.17 |
| ProtT5 (XL-U50) | 0.56 ± 0.20 | 0.49 ± 0.14 | 0.59 ± 0.21 | 0.55 ± 0.20 |

On the RemoteFoldSet benchmark, the ESM family and CARP exhibit the strongest structural awareness, with SA scores typically around 0.60, and CARP achieving up to 0.70 on beta structures. The ProtTrans models (ProtBert and ProtT5) form the second tier, reaching scores of approximately 0.56. UniRep performs less effectively, with a score of 0.52, while the ProGen2 models show the weakest structural awareness, with scores ranging from 0.33 to 0.45.

In comparisons within the same model family, we observe that for ProGen2, increasing the parameter count from 764M (base) to 6.4B (xlarge) does not lead to an improvement in SA score, and even results in a decrease for the xlarge model. In contrast, for ESM-2, scaling from 650M to 3B yields an improvement in SA score, although the score decreases again at 15B. These results suggest that the ability to capture structural information does not necessarily increase with the size of pLMs. The compatibility between the model architecture and the training objective might play a critical role in determining whether scaling translates into structural awareness.

A closer look at the differences between ESM-2 and ProGen2 helps illustrate this point (Table 4). ESM-2 is trained with a masked language modeling (MLM) objective, which leverages bidirectional context and may be more conducive to learning embeddings that capture comprehensive structural information through richer contextual representation. This characteristic also contributes to the strong performance of ESMFold, which is built upon ESM-2 embeddings. In contrast, ProGen2 adopts a GPT-style autoregressive architecture trained for next-token prediction, which is more optimized for sequence generation than for producing globally informative representations.

An interesting observation is that UniRep performs particularly well on alpha structures. Given that UniRep has a very simple architecture consisting of only a single-layer mLSTM, it is capable of capturing local information that aligns with the primarily local nature of helices. This explains its relative strength on alpha structures but weaker performance on more complex classes. However, its superior scores in the Mainly Alpha category also reveal a limitation of the SA metric: UniRep may be capturing structural patterns, but these likely correspond to shared local features that are present across different alpha proteins, and therefore do not enable reliable distinction between them. Consistent with this, UniRep exhibits a substantially worse SA distance ratio (Appendix A.3) than other models in the Mainly Alpha category in Table 5, suggesting that its structural awareness may be largely due to inductive biases of the architecture rather than truly informative learned representations.

In the case of CARP, its optimal SA distance ratio in the Mainly Beta category of Table 5 suggests that the high SA score is not merely indicative of general structural awareness, but reflects precise and discriminative information. One possible explanation for CARP’s strong performance on beta structures is its ByteNet-based dilated convolution architecture, which may more stably preserve the spatial pairing patterns required for beta-sheet formation. While dilated convolutions are theoretically less capable than self-attention at modeling arbitrary long-range dependencies, their progressively expanding receptive fields could confer an advantage in maintaining such pairing patterns, potentially leading to better structural awareness.

### 4.2 Validating the structural awareness metric

To ensure that the proposed SA score genuinely reflects fold-level structural information rather than artifacts or dataset biases, we conduct a control experiment using shuffled embeddings. Although mean-centering removes global biases, we still need to account for two potential sources of error: residual confounding factors, such as dominant principal components in the embedding space, and possible limitations of the SA scoring procedure itself. To this end, for each model, we randomly permuted the 1,440 embeddings, completely breaking the original alignment between sequences and embeddings. These shuffled embeddings were randomly grouped into 90 new sets of 16 sequences each, and SA scores were computed using the same methodology. The corresponding results are shown in Figure 2.

Panels (A) and (B) display the SA scores obtained from the original embeddings of two representative and widely used models: ESM-2 (650M) and ProtT5 (XL-U50), respectively. Each of the 90 bars corresponds to one of the protein sequence sets in the RemoteFoldSet dataset. The SA scores range from approximately 0.1 to 1.0, with most values clustering around 0.6, in agreement with the averages reported in Table 1. Moreover, the distribution patterns of the two models exhibit similar trends, indicating that both models capture comparable levels of structural information across different protein sets. For the shuffled embeddings in panels (C) and (D), the SA scores drop sharply, all falling within a narrow range around zero (-0.1 to 0.1), and the distributions show no discernible patterns. These near-zero values and random patterns confirm that the embeddings no longer exhibit consistent representational clustering, and that the SA score does not mistakenly capture similarity driven by noise.

### 4.3 Layer-wise structural awareness analysis

To investigate how structural information is captured across different depths of pLMs, we perform a layer-wise evaluation on ESM-2 and ProtT5. For each Transformer layer, we compute the structural awareness score and visualize its evolution, revealing where structural patterns emerge and how they change throughout the network.

From the layer-wise structural awareness curves shown in Figure 3, we observe a consistent pattern across different models: the scores are moderately high in the early layers, drop in the lower-middle layers, and then steadily increase in the mid-to-late layers before declining again near the final layers. This trend suggests that the early layers capture basic structural cues by focusing on local sequence patterns and token-level representations. As the model progresses, the structural awareness temporarily decreases, possibly due to the integration of more contextual or semantic information, which may obscure direct structural signals. In the mid layers, structural features appear to be reinforced, leading to a rise in structural awareness that stabilizes into a plateau. This phase likely reflects the model’s ability to internalize fold-level patterns. However, in the final layers, the structural awareness declines again, potentially due to the representations being adapted or optimized for downstream objectives (e.g., MLM), which may deprioritize fold-specific features.

**Figure 3:**
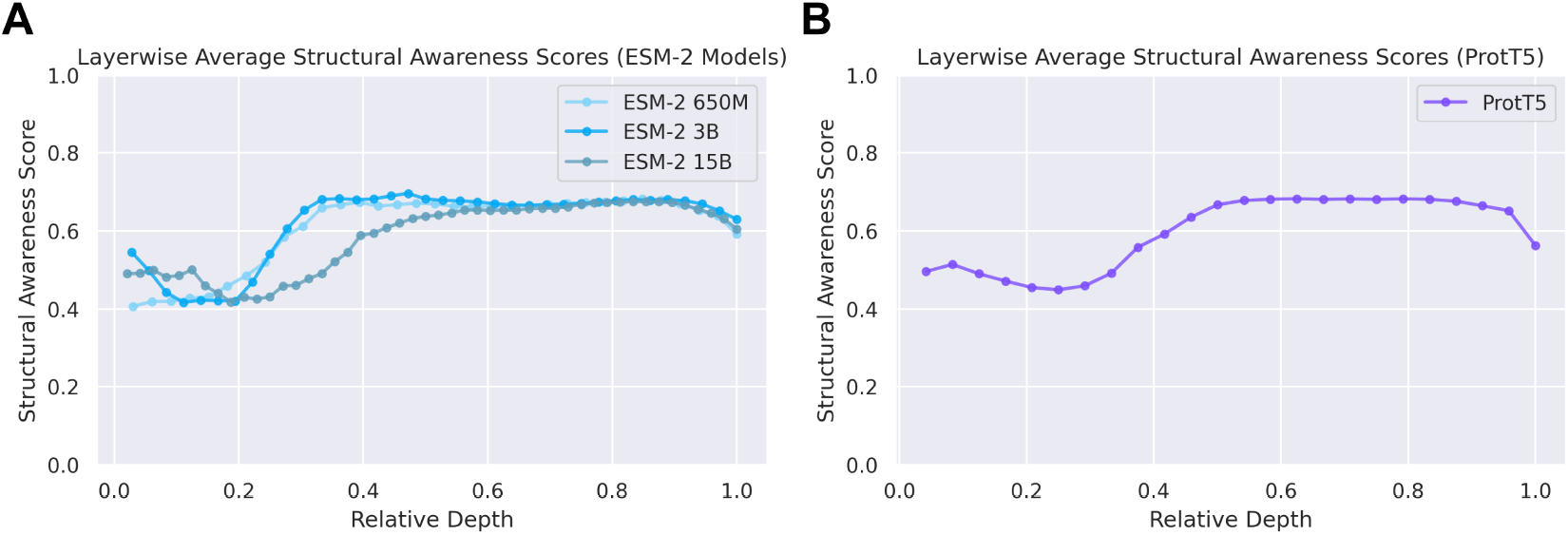
Layer-wise structural awareness of pLMs. **(A)** Structural awareness scores across transformer layers for three ESM-2 models of varying scales (650M, 3B, 15B parameters). **(B)** Layer-wise structural awareness for ProtT5 (XL-U50).

Within the ESM-2 model family, we observe that the early to mid layer drop in structural awareness becomes more pronounced as the model size increases. The smallest model (650M) exhibits a relatively flat trajectory, while the 3B and 15B variants show a clear dip followed by a recovery. Interestingly, ProtT5, which has approximately 3 billion parameters, shows a similar trend to ESM-2 (3B), further supporting the observation that the magnitude of this drop is closely related to model capacity. This suggests that larger models tend to perform more complex internal reorganization of representations between layers, which may temporarily reduce the prominence of low-level structural signals before they are recovered in later layers.

### 4.4 Visualization of embedding representations

In addition to quantitative evaluation, we also conducted t-SNE visualization of the learned embeddings to qualitatively compare the performance of different pLMs. Specifically, we applied t-SNE directly to the 1,440 protein sequence embeddings obtained from each model, using a perplexity of 30 and a learning rate of 200. The models included in the comparison are ProGen2 (base), UniRep, CARP (640M), ESM-2 (650M), and ProtT5 (XL-U50). As a baseline, we also included representations obtained by mean-pooling over one-hot encoded sequences. The t-SNE projections are presented in Figure 4.

**Figure 4:**
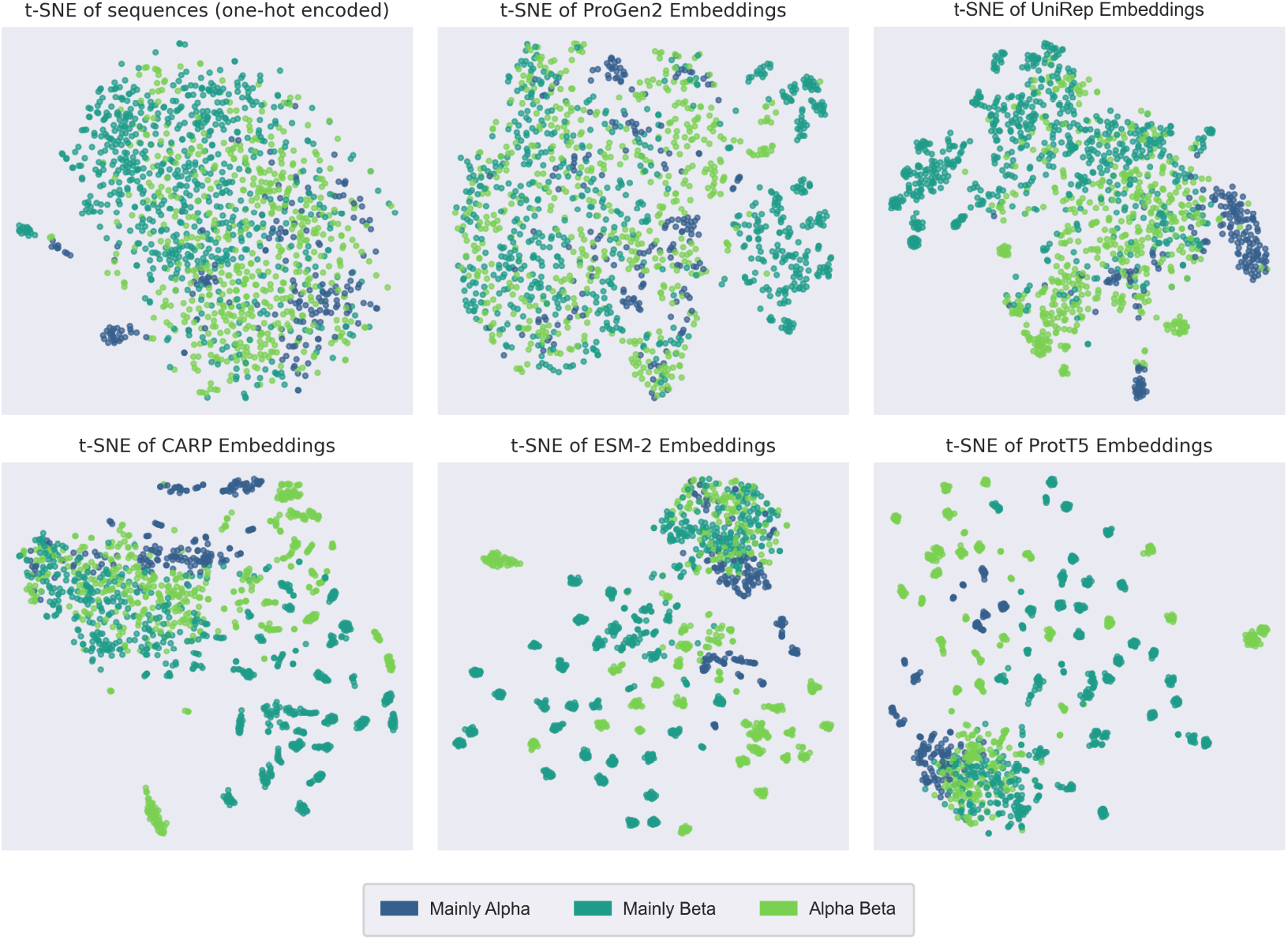
t-SNE visualizations of protein sequence embeddings. Each plot shows the t-SNE projection of 1,440 sequences represented by different models, with points colored based on the class-level labels from CATH. The top row (from left to right) corresponds to the one-hot encoded baseline, ProGen2 (base) and UniRep, while the bottom row shows CARP (640M), ESM-2 (650M), and ProtT5 (XL-U50).

The one-hot encoded baseline exhibits an unstructured distribution, reflecting its limited discriminative capacity. ProGen2 shows a modest refinement over the baseline, with a few noticeable clusters emerging in the upper-right region, although most points remain dispersed without clear organization. Compared to ProGen2, UniRep embeddings demonstrate a slightly more structured distribution, with clearer groupings and several well-defined clusters. In contrast, the t-SNE plots of CARP, ESM-2, and ProtT5 embeddings reveal numerous compact clusters separated by class, indicating a stronger ability to capture structural similarity. Among them, ESM-2 and ProtT5 exhibit the most distinct patterns, characterized by a higher number of tight clusters with well-defined intra-cluster cohesion and inter-cluster separation.

These visualization results are consistent with the quantitative trend of the SA score in Table 1. ProGen2 exhibits the lowest score and weakest clustering, followed by UniRep, while CARP, ESM-2, and ProtT5 all demonstrate substantially better performance and are comparably strong. This alignment further supports the SA score as a faithful and model-agnostic indicator of how effectively different pLMs capture structural information from sequences.

## 5 Conclusion

In this work, we introduce RemoteFoldSet, a new dataset and evaluation framework designed to quantify the intrinsic structural knowledge encoded by pLMs. By benchmarking a wide range of pLMs, we observe that ESM-2 (3B) exhibits the strongest capacity to encode structural information. CARP also demonstrates strong performance, particularly on beta proteins, which may stem from its unique ByteNet-based dilated convolution architecture. Interestingly, model size alone does not directly correlate with structural awareness, as architectural design and training objectives appear to play a more critical role. For example, BERT-style model embeddings are more structurally informative compared to their GPT-style counterparts. Additionally, layer-wise analysis reveals a non-monotonic trend in fold-level structural encoding, with an initial dip followed by a steady increase across layers. This may indicate that early to mid layers in large models perform a degree of representational integration, temporarily suppressing explicit structural signals before they are reinforced in deeper layers.

More broadly, our results highlight the importance of evaluating the intrinsic representational quality of pLMs. While widely-used benchmarks such as TAPE rely on supervised probing tasks that approximate structural knowledge through downstream performance, task-specific approximation is fundamentally distinct from task-free intrinsic evaluation. By explicitly decoupling sequence similarity from structural similarity in dataset design, RemoteFoldSet enables direct assessment of the structural information encoded in protein embeddings. This design allows the SA score to be computed in a simple yet meaningful way, and also makes structural patterns readily observable through unsupervised visualization methods such as t-SNE, which is not possible with prior evaluation frameworks.

RemoteFoldSet also provides substantial practical advantages, enabled by its simple, training-free evaluation procedure. Assessing structural awareness with SA scores requires only a single forward pass per sequence, making it orders of magnitude faster than supervised probing approaches, particularly for layer-wise analysis across large models. As pLMs continue to scale in size and application, we expect RemoteFoldSet to serve as a lightweight, interpretable, and scalable diagnostic tool for analyzing their structural representational capacity.

## Acknowledgements

We thank the HHMI Hanna Gray Fellowship Grant #17718 and the Powell-Bundle Fellowship for funding, and the San Diego Supercomputer Center (SDSC) and UCSD Physics Computing Facility for computational resources.

## Code availability

The curated dataset and source code of this work are available at https://github.com/ZinniaMa/RemoteFoldSet.

## A Supplementary Methods

### A.1 Dataset generation details

We selected 168 domains from the CATH database, aiming for an approximately even coverage across 43 architectures. To reduce redundancy, no two domains were taken from the same superfamily. For sequence generation, we provided the CATH structures directly to ProteinMPNN. In cases where the input structures contained unresolved regions, ProteinMPNN left ‘X’ characters in the generated sequences as placeholders. We retained these ‘X’ positions because they implicitly reflect spatial constraints that may be useful for downstream modeling.

The ProteinMPNN-generated sequences, including the ‘X’ placeholders, were then submitted to AlphaFold3 for structure prediction. Because AlphaFold3 attempts to generate complete structures even when ‘X’ placeholders are present, the resulting predictions contained no missing regions. For evaluation, we computed TM-scores over residues corresponding to regions with resolved structures in the CATH templates, thereby excluding originally missing segments. This ensures that the comparison reflects only the experimentally validated parts of the structures. Meanwhile, the pTM scores were taken directly from AlphaFold3’s output summary files.

For each domain, we generate 160 sequences using ProteinMPNN and retain only those satisfying both pTM > 0.8 and TM-score > 0.8. If fewer than 16 sequences pass these thresholds, the corresponding domain is then discarded. Otherwise, we perform a greedy search to select a diverse subset of 16 sequences. The selection begins with a single seed sequence, and sequences are added iteratively by choosing the one that maximizes the diversity of the current set. This process is repeated with each candidate sequence as the initial seed, and the final selection is the one that achieves the highest overall diversity. When computing diversity, we considered only the non-’X’ positions in the sequences.

When providing sequences to pLMs, we preserved the ‘X’ positions to enable the models to capture information about the lengths and positions of missing regions. After obtaining residue-level embeddings for the full sequences, we removed embeddings corresponding to the ‘X’ positions and performed mean pooling over the remaining embeddings to produce the final protein-level representations. This step ensures that the final representations reflect only biologically meaningful residues, avoiding artifacts that could otherwise bias the calculation of the SA score.

### A.2 Hyperparameter selection

To evaluate the impact of the sampling module’s temperature on the stochasticity of sequence generation in inverse folding, we systematically varied the temperature and analyzed its effect on several evaluation metrics. One domain was selected from each of the CATH architectures, and ProteinMPNN was used to generate sequences conditioned on these structures. For each temperature setting (1.0, 1.5, and 2.0), we sampled 160 sequences per structure and evaluated them in terms of Recovery, Diversity, Root Mean-Squared Deviation (RMSD), and TM-score.

As shown in Table 2, increasing the sampling temperature significantly affects structural fidelity. At *T* = 1.5, the RMSD reaches 12 Å while the TM-score drops to 0.5, indicating that most generated sequences fail to maintain the original fold. At *T* = 2.0, both metrics further deteriorate, suggesting that temperatures of 1.5 or higher are not suitable for our goal of generating fold-consistent sequences. In contrast, at *T* = 1.0, the TM-score remains around 0.8, which is acceptable for our use case. Although the diversity at this temperature is relatively low (0.72), it can be slightly improved using post hoc greedy search strategies. If the temperature is set even lower, the fold consistency may improve further, but the resulting diversity would fall below the level required for our dataset construction goals. Considering the trade-off between diversity and refoldability, we selected *T* = 1.0 for sequence sampling during dataset construction.

**Table 2:**
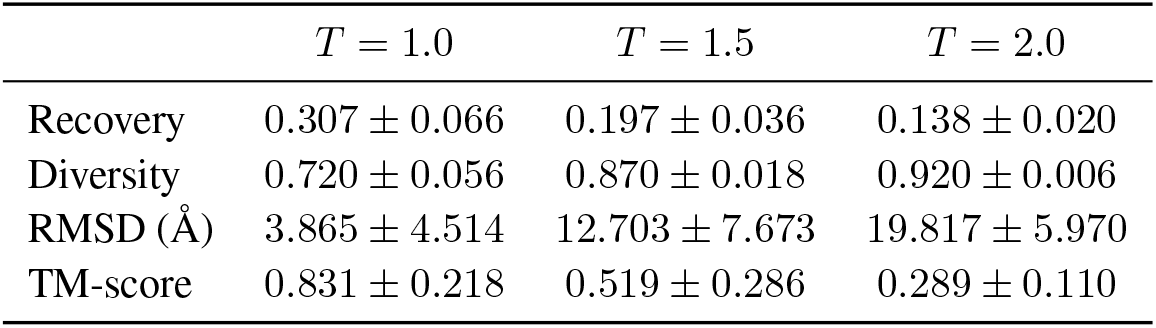
Performance across different temperature (*T*) values.

| | $T = 1.0$ | $T = 1.5$ | $T = 2.0$ |
| --- | --- | --- | --- |
| Recovery | $0.307 \pm 0.066$ | $0.197 \pm 0.036$ | $0.138 \pm 0.020$ |
| Diversity | $0.720 \pm 0.056$ | $0.870 \pm 0.018$ | $0.920 \pm 0.006$ |
| RMSD (Å) | $3.865 \pm 4.514$ | $12.703 \pm 7.673$ | $19.817 \pm 5.970$ |
| TM-score | $0.831 \pm 0.218$ | $0.519 \pm 0.286$ | $0.289 \pm 0.110$ |

For the final version of the dataset, we selected 168 domains, corresponding to approximately 5 domains per architecture. If an architecture contained fewer domains, we included as many as were available, aiming for broad structural coverage. For each domain, we sampled 160 sequences using a temperature of *T* = 1.0, resulting in a total of 26,880 sequences. To ensure reasonable refoldability, we measured how many of the generated sequences passed certain thresholds defined by combinations of pTM and TM-score, as summarized in Table 3. Notably, pTM has a greater impact on the number of sequences passing the thresholds compared to TM-score. When applying a strict threshold of pTM *>* 0.9, only about 6% of sequences passed, which was too low and left multiple domains with no qualifying sequences, compromising structural coverage. In contrast, with pTM *>* 0.8, approximately 36% of sequences passed, which we consider an acceptable balance. For TM-score, we chose a threshold of 0.8, striking a compromise between better alignment and sufficient sequence retention, and used it in combination with the pTM threshold.

**Table 3:** Number of samples meeting joint thresholds on pTM and TM-score.

|  | TM-score > 0.7 | TM-score > 0.8 | TM-score > 0.9 |
| --- | --- | --- | --- |
| pTM > 0.7 | 14813 / 26880 | 14578 / 26880 | 12315 / 26880 |
| pTM > 0.8 | 9902 / 26880 | 9893 / 26880 | 9357 / 26880 |
| pTM > 0.9 | 1708 / 26880 | 1708 / 26880 | 1708 / 26880 |

### A.3 Metrics

#### TM-score

The template modeling score (TM-score) is a length-normalized metric for evaluating global structural similarity between two protein structures. It ranges from 0 to 1, with scores above 0.5 indicating roughly correct topology[16][17].

#### pTM

The predicted template modeling (pTM) score is a confidence metric from AlphaFold 3 that estimates the TM-score, which reflects the accuracy of the predicted protein fold. Higher pTM scores indicate that the predicted structure should be more similar to the true structure[7].

#### Diversity

Let *m* be the number of generated sequences, each of length *n*. Let *r*_*i,k*_ denote the residue at position *k* in the *i*-th sequence, where 1 ≤ *i* ≤ *m* and 1 ≤ *k* ≤ *n*. We compute diversity as the average pairwise mismatch ratio between all sequence pairs:

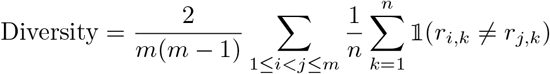

#### Recovery

When using the inverse folding to generate alternative sequences for the given structure, the sequence recovery rate measures the identity between the generated sequences and the reference sequence[6][18].

#### SA distance ratio

For each structural set 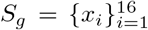, the model *f* produces sequence embeddings *z*_*i*_ = *f* (*x*_*i*_), which are first mean-centered to remove global bias. Let *G* denote the total number of sets, and define the group mean embedding as *µ*_*g*_. Using the distance as one minus cosine similarity, we compute the average intra- and inter-group distances, and take their ratio as the SA distance ratio:

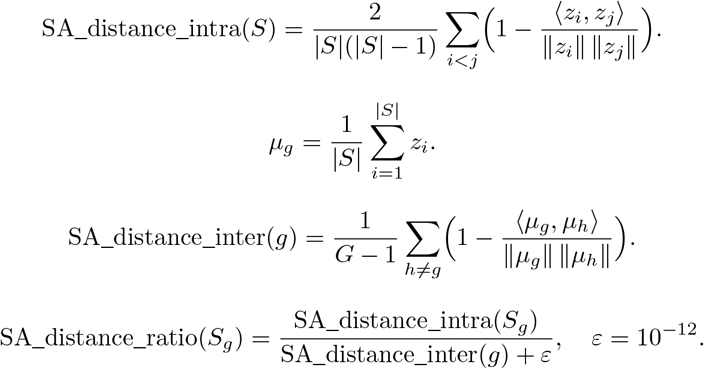

## B Dataset Details

### B.1 Sequence diversity

**Figure 5:**
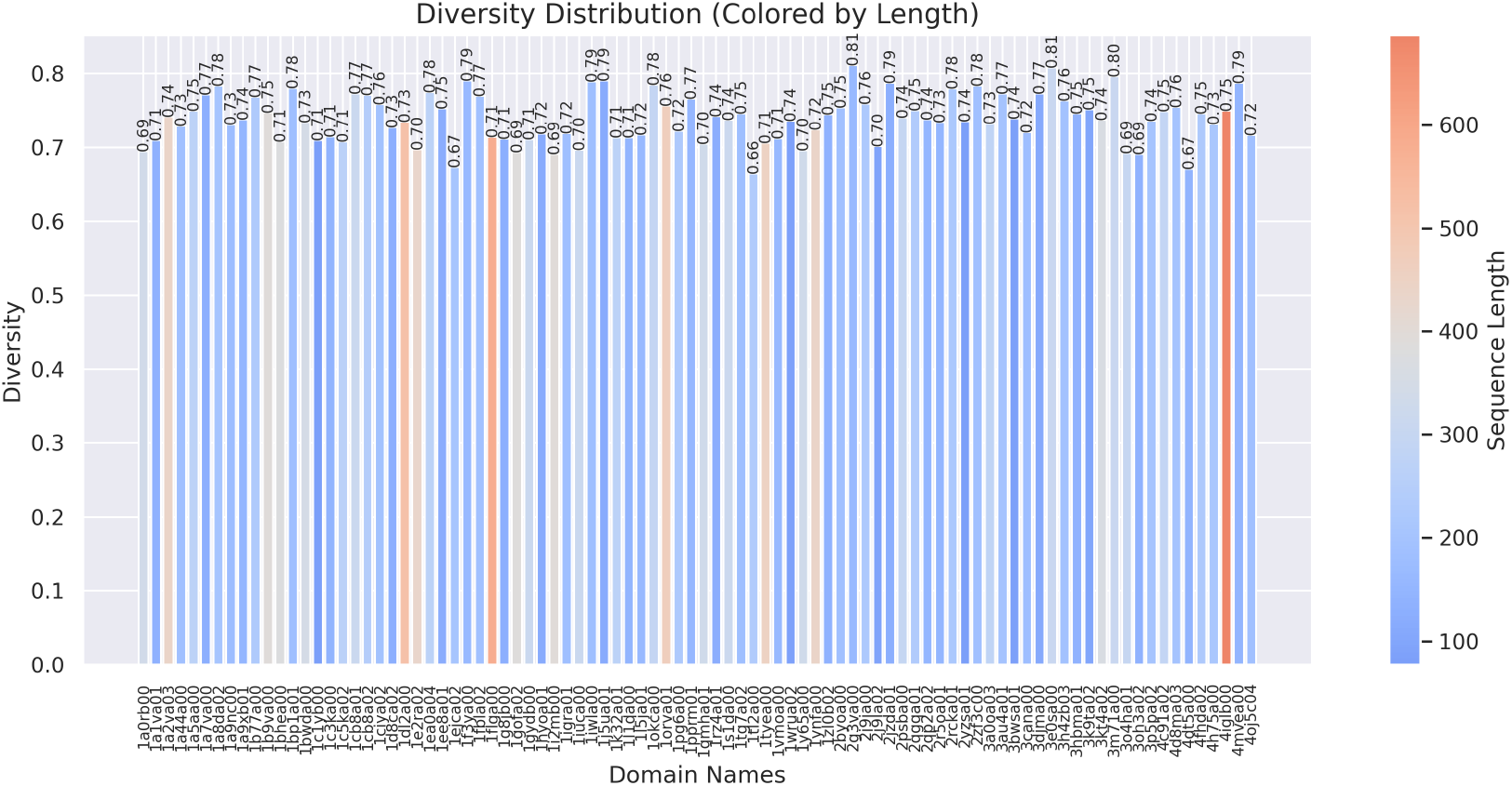
Diversity distribution across all protein sets. Colored by sequence length. Each set is labeled by its domain name.

### B.2 Structural similarity

**Figure 6:**
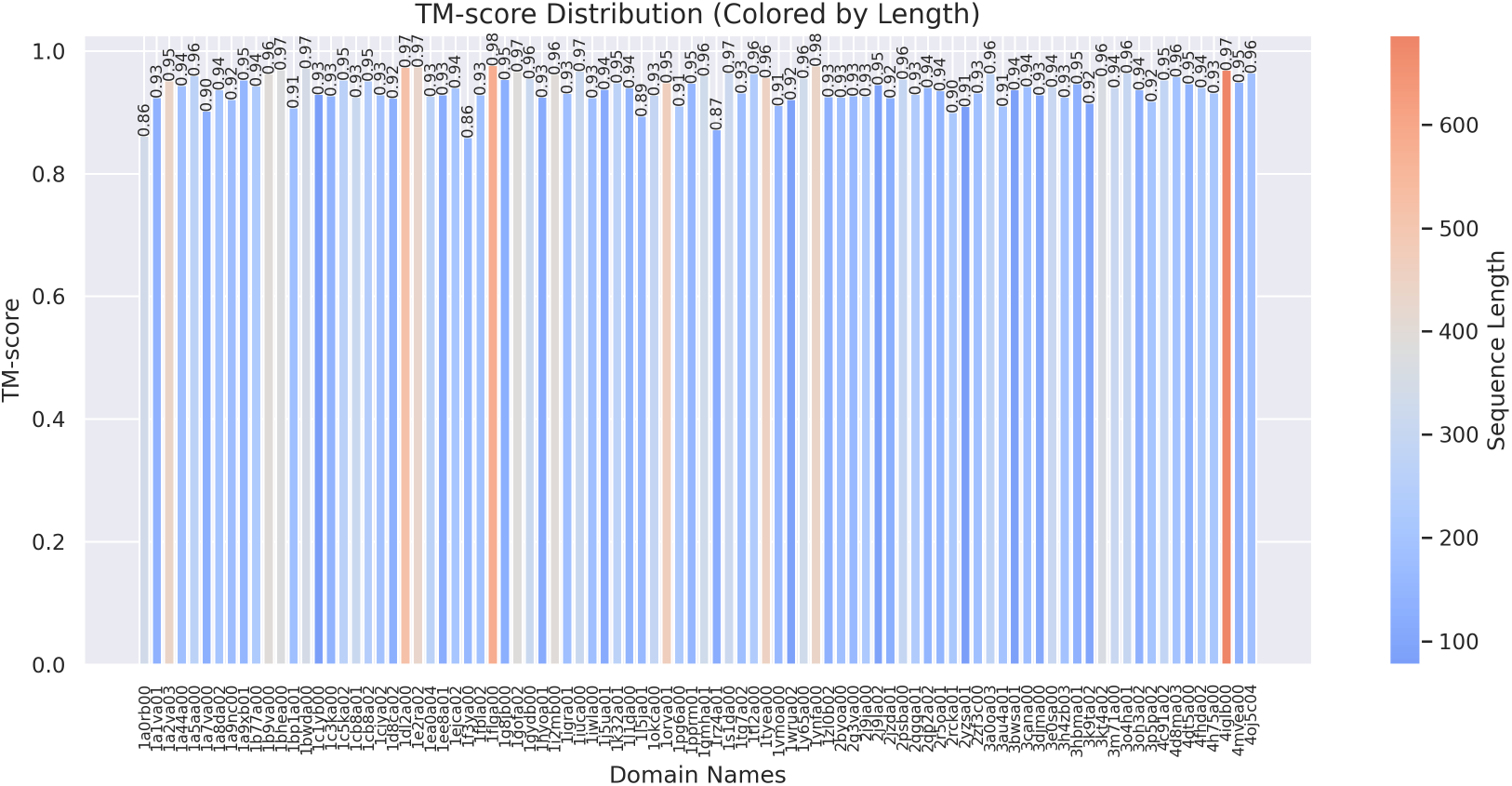
TM-score distribution across all protein sets. Visualization follows the same format as Figure 5.

## C Evaluated Protein Language Models

**Table 4:** Comparison of pLMs.

| Model | Architecture | Objective | Size | Layers | Embedding |
| --- | --- | --- | --- | --- | --- |
| UniRep | mLSTM | Next-token | 18M | 1 | 1900 |
| ProGen2 (base) | GPT-style | Next-token | 764M | 27 | 1536 |
| ProGen2 (large) | GPT-style | Next-token | 2.7B | 32 | 2560 |
| ProGen2 (xlarge) | GPT-style | Next-token | 6.4B | 32 | 4096 |
| CARP (640M) | ByteNet-style | MLM | 640M | 30 | 1280 |
| ESM-1b | BERT-style | MLM | 650M | 33 | 1280 |
| ESM-2 (650M) | BERT-style | MLM | 650M | 33 | 1280 |
| ESM-2 (3B) | BERT-style | MLM | 3B | 36 | 2560 |
| ESM-2 (15B) | BERT-style | MLM | 15B | 48 | 5120 |
| ProtBert | BERT-style | MLM | 420M | 30 | 1024 |
| ProtT5 (XL-U50) | T5-style | Span MLM | 3B | 24 (enc) / 24 (dec) | 1024 |

## D Supplementary Experiments

### D.1 Benchmarking protein language models using SA distance ratio

**Table 5:** Comparison of the SA distance ratio of different pLMs (lower is better).

| Models | All domains | Mainly Alpha | Mainly Beta | Alpha Beta |
| --- | --- | --- | --- | --- |
| UniRep | $0.50 \pm 0.21$ | $1.01 \pm 0.39$ | $0.52 \pm 0.26$ | $0.73 \pm 0.22$ |
| ProGen2 (base) | $0.57 \pm 0.23$ | **$0.69 \pm 0.26$** | $0.52 \pm 0.26$ | $0.69 \pm 0.21$ |
| ProGen2 (large) | $0.57 \pm 0.20$ | $0.74 \pm 0.21$ | $0.55 \pm 0.23$ | $0.75 \pm 0.16$ |
| ProGen2 (xlarge) | $0.68 \pm 0.26$ | $0.92 \pm 0.48$ | $0.63 \pm 0.25$ | $0.86 \pm 0.24$ |
| CARP (640M) | $0.40 \pm 0.29$ | $0.77 \pm 0.34$ | **$0.30 \pm 0.27$** | $0.51 \pm 0.32$ |
| ESM-1b | $0.41 \pm 0.22$ | $0.71 \pm 0.31$ | $0.37 \pm 0.23$ | $0.47 \pm 0.22$ |
| ESM-2 (650M) | $0.41 \pm 0.22$ | $0.76 \pm 0.35$ | $0.36 \pm 0.21$ | $0.45 \pm 0.22$ |
| ESM-2 (3B) | **$0.37 \pm 0.20$** | $0.71 \pm 0.32$ | $0.33 \pm 0.20$ | **$0.40 \pm 0.19$** |
| ESM-2 (15B) | $0.39 \pm 0.19$ | $0.70 \pm 0.31$ | $0.35 \pm 0.20$ | $0.43 \pm 0.18$ |
| ProtBert | $0.46 \pm 0.20$ | $0.71 \pm 0.21$ | $0.46 \pm 0.24$ | $0.54 \pm 0.19$ |
| ProtT5 (XL-U50) | $0.44 \pm 0.21$ | $0.70 \pm 0.24$ | $0.42 \pm 0.22$ | $0.46 \pm 0.21$ |

